# Structure and ion-release mechanism of P_IB-4_-type ATPases

**DOI:** 10.1101/2021.09.01.458532

**Authors:** Christina Grønberg, Qiaoxia Hu, Dhani Ram Mahato, Elena Longhin, Nina Salustros, Annette Duelli, Jonas Eriksson, Komal Umashankar Rao, Domhnall Iain Henderson, Gabriele Meloni, Magnus Andersson, Tristan Croll, Gabriela Godaly, Kaituo Wang, Pontus Gourdon

## Abstract

Transition metals, such as zinc, are essential micronutrients in all organisms, but also highly toxic in excessive amounts. Heavy-metal transporting P-type (P_IB_) ATPases are crucial for homeostasis, conferring cellular detoxification and redistribution through transport of these ions across cellular membranes. No structural information is available for the P_IB-4_-ATPases, the subclass with the broadest cargo scope, and hence even their topology remains elusive. Here we present structures and complementary functional analyses of an archetypal P_IB-4_-ATPases, sCoaT from Sulfitobacter sp. NAS14-1. The data disclose the architecture, devoid of classical so-called heavy metal binding domains, and provides fundamentally new insights into the mechanism and diversity of heavy-metal transporters. We reveal several novel P-type ATPase features, including a dual role in heavy-metal release, and as an internal counter ion, of an invariant, central histidine. We also establish that the turn-over of P_IB_-ATPases is potassium independent, contrasting to many other P-type ATPases. Combined with new inhibitory compounds, our results open up for efforts in e.g. drug discovery, since P_IB-4_-ATPases function as virulence factors in many pathogens.

## Introduction

The ability to adapt to environmental changes in heavy metal levels is paramount for all cells, as these elements are essential for a range of cellular processes and yet toxic at elevated concentrations ^(*1, 2*)^. Transition metal transporting P-type (P_IB_) ATPase proteins are critical for cellular heavy metal homeostasis, providing efflux of e.g. copper, zinc and cobalt from the intracellular milieu. Indeed, malfunctioning of the human P_IB_-members, ATP7A and ATP7B, cause the fatal neurological Menkes disease and Wilson disease ^(*3, 4*)^. The P_IB_-ATPases belong to the P-type ATPase superfamily of integral membrane proteins, which exploit energy from ATP hydrolysis for transport of cargo across cellular membranes. These proteins share an overall mechanism described by the so-called Post-Albers cycle ^(*5, 6*)^, as established by decades of structural and functional investigations of primarily Ca^2+^-, Na^+^/K^+^- and H^+^-specific P-type ATPases^(*7-15*)^. In summary, four cornerstone states, E1-E1P-E2P-E2, provide alternating access and affinity for the transported ions (and counter-ions, if present). Inward facing (e.g. cytosolic) E1 and outward facing (e.g. extracellular) E2P conformations are coupled to ATP-dependent phosphorylation (yielding ion-occluded E1P) and dephosphorylation (to occluded E2) of an invariant catalytical aspartate, respectively.

P_IB_-ATPases are subdivided into groups based on conserved sequence motifs and the selectivity towards transported transition metal ions^(*16-19*)^. Whereas Cu^+^- and Zn^2+^-transporting P_IB-1_ and P_IB-2_ ATPases are relatively well-characterized, little is known regarding the P_IB-4_ proteins, which comprise some of the simplest and shortest proteins within the entire P-type ATPase superfamily^(*16*)^. They are present in plants, archaea and prokaryotes, and have been assigned a role as virulence factors in pathogens, as e.g. the P_IB-4_-ATPase MtCtpD is required for tuberculosis infections^(*20, 21*)^, and therefore represent attractive targets for novel antibiotics.

The P_IB-4_-ATPases are classically referred to as cobalt transporters. However, the metal specificity of the P_IB-4_ ATPases remain elusive as some members have a confirmed cobalt-specificity, while others seemingly have broader or altered ion-transport profiles, transporting also ions such as Zn^2+^, Ni^2+^, Cu^+^ and even Ca^2+ (*18, 22-25*)^. Thus, the P_IB-4_-ATPases appear to have the widest scope of transported ions of the P_IB_-ATPases, and it is possible that further sub-classification principles and sequence motifs will be identified. Due to the broad ion transport range, they have been proposed to serve as multifunctional emergency pumps that can be exploited under extreme environmental stress to maintain heavy metal homeostasis^(*26*)^.

Hitherto, the available high-resolution structural information of full-length P_IB_-ATPases is limited to two structures each of ion-free conformations of the Cu^+^-transporting P_IB-1_-ATPase from *Legionella pneumophila* (LpCopA)^(*27, 28*)^, and the Zn^2+^-transporting P_IB-2_-ATPase from *Shigella sonnei* (SsZntA)^(*28*)^. Thus, the principal architecture of the P_IB-4_-ATPases remains debated, as sequence analyses have proposed different topologies for the N-terminus: with or without i) the so-called called heavy metal binding domains (HMBDs), and ii) the first two transmembrane helices, MA and MB^(*16, 29-31*)^, which both are present in other P_IB_-ATPases (**Supplementary Figure 1a**). These represent structural features that have been suggested to be important for ion-uptake and/or regulation in other P_IB_-ATPases^(*27, 28, 32, 33*)^, raising questions if these levels of protein control are absent or replaced in the P_IB-4_ group. In addition, despite a shared overall architecture, the P_IB-1_ and P_IB-2_ structures suggested significantly different types of entry and exit pathways, hinting at unique translocation mechanisms for each P_IB_ group^(*34*)^. However, it remains unknown if similar molecular adaptions have taken place in P_IB-4_-ATPases to handle the unique array of cargos. To address these fundamental questions, we determined structures of a P_IB-4_-ATPase in different states and validated our findings using *in vitro* functional characterization and molecular dynamics simulations.

## Results & Discussion

### Metal specificity

We employed the established P_IB-4_ model sCoaT (UniProt ID A3T2G5) to shed further light on the structure and mechanism of the entire P_IB-4_-class. As the metal ion specificity of the P_IB_-4-ATPases is known to be wide, the ATPase activity was assessed *in vitro* in lipid-detergent solution, in the presence of a range of different heavy metals. The protein exhibited clear Zn^2+^ and Cd^2+^ dependent ATPase activity, while Co^2+^ only stimulated ATP-hydrolysis at high ion concentrations. This is in partial agreement with the ion range profile previously reported for sCoaT, as higher Co^2+^ sensitivity has been detected using a different functional assay and different experimental conditions^(*18*)^(**Supplementary Figure 2**). However, it cannot be excluded that Co^2+^, rather than Zn^2+^, is the preferred cargo *in vivo* as the relative intracellular availability of Co^2+^ is more than three orders of magnitude higher than that of Zn^2+^ in certain bacterial cells^(*35*)^.

### Structure determination

We determined structures of sCoaT in metal-free conditions supplemented with two different phosphate analogues, BeF_3-_ and AlF_4-_, respectively, which previously have been exploited to stabilize E2 reaction intermediates of the transport cycle. The structures were determined at 3.1 Å and 3.2 Å resolution, using molecular replacement as phasing method and SsZntA as search model, and the final models yielded R/R_free_ of 24.4/26.8 and 21.8/25.5 (**Supplementary Table 1**). The two crystal forms were obtained using the HiLiDe method (crystallization in the presence of high concentrations of detergent and lipids)^(*36*)^. Surprisingly however, the crystal packing for both structures reveal only minor contacts between adjacent membrane-spanning regions, which are critical for the crystals obtained of most other P-type ATPase proteins (**Supplementary Figure 3**). Hence, some crystal forming interactions likely take place through lipid-detergent molecules. To our knowledge, this is the first time that type I crystals with unrestrained transmembrane domains are reported, but a consequence is that peripheral parts of the membrane domain are less well-resolved (**Supplementary Figure 4**). While this caused difficulty in modelling some transmembrane helices, satisfying solutions were found with the aid of the software ISOLDE^(*37*)^ due to its use of AMBER forcefield which helped to maintain physical sensibility in the lowest-resolution regions.

### Overall structure, without classical HMBD

Examination of the structures reveals that the P_IB-4_-ATPase architecture is reminiscent to that of other P-type ATPases, with three cytosolic domains, A (actuator), N (nucleotide-binding) and P (phosphorylation), as well as a membrane spanning M-domain (**Figure 1a**). Furthermore, the core of the soluble portions, including the nucleotide binding pocket and catalytic phosphorylation site at D369, are well-conserved.

**Figure 1.**
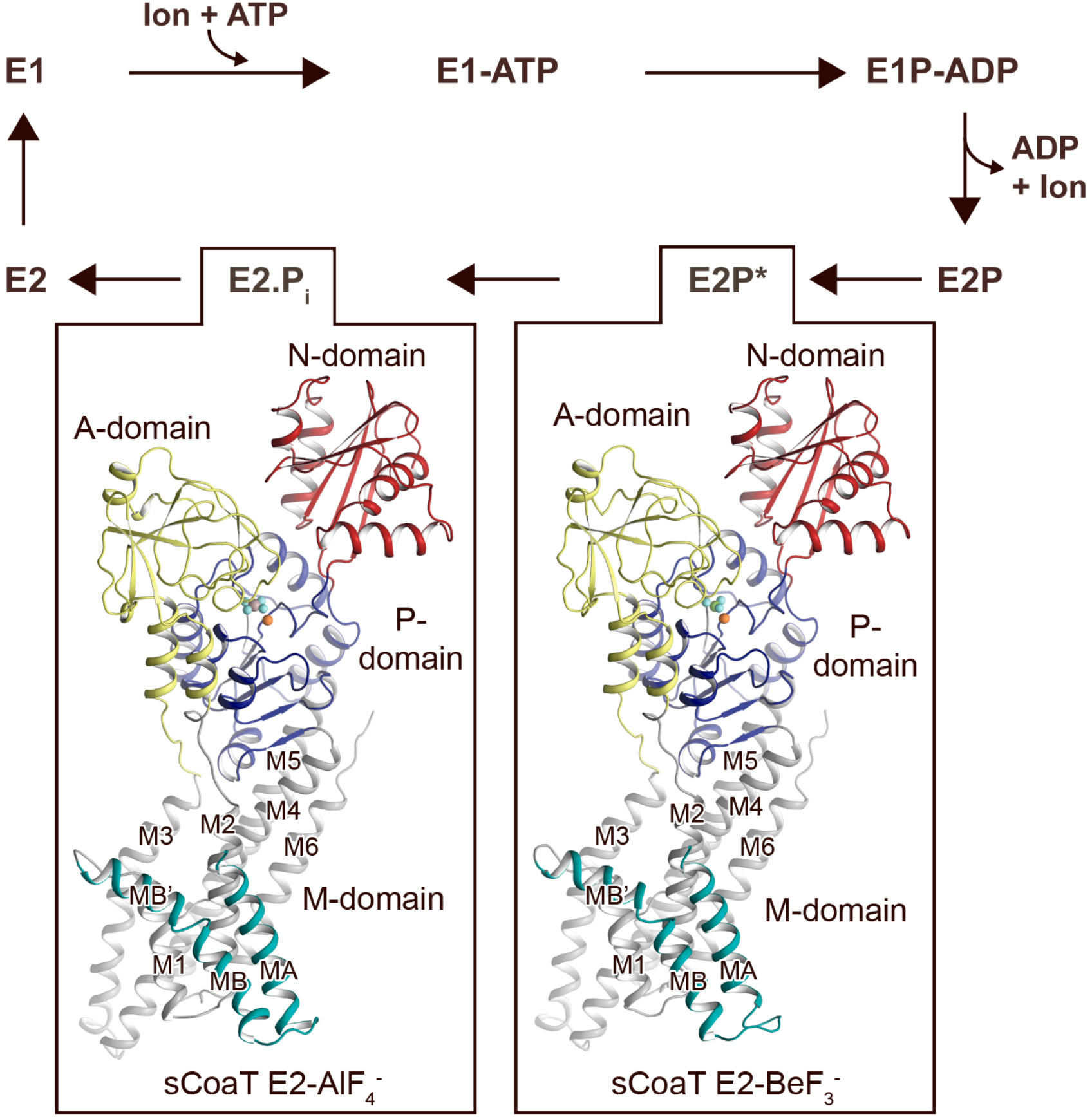
Overall architecture and reaction cycle. The sCoaT structures reveal that P_IB-4_-ATPases comprise soluble A-, P- and N-domains, shown in yellow, blue and red, respectively, as well as a transmembrane domain with eight helices: MA and MB, in cyan, and M1-M6, in grey, and that the P_IB-4_-topology lacks classical so-called heavy metal binding domain. The transport mechanism of P-type ATPases depends on ATP-dependent phosphorylation and auto-dephosphorylation, and include four principal conformations, E1, E1P, E2P and E2, where P denote phosphorylation. The determined structures are trapped in two transition states following ion-release – an occluded late E2P (E2P^*^) and an occluded transition state of dephosphorylation, E2.P_i._

The topology of P_IB-4_-ATPases has been a conundrum as sequence analyses have proposed different arrangements, with variable number of transmembrane segments and different sizes of the N-termini^(*16, 27, 28, 30, 31, 38*)^. However, our data unambiguously demonstrate that P_IB-4_-ATPases possess eight transmembrane helices, MA and MB followed by M1-M6. As previously observed for P_IB-1_- and P_IB-2_-ATPases, MB is kinked by a conserved Gly-Gly motif (G82 and G83) forming an amphipathic ‘platform’, MB’, immediately prior to M1, see further below (**Supplementary Figure 4a-b)**.

Are then HMBDs present in P_IB-4_-ATPases as in the other P_IB_ subclasses? As only the first 47 residues remain un-modelled in the final structures (**Supplementary Table 1**), it is clear that many P_IB-4_-ATPases including sCoaT are lacking a classical HMBD ferredoxin-like fold (typically 70 residues long). In agreement with this observation, the cysteine pair (CGIC in the sequence) in the N-terminus of sCoaT is rather positioned in MA, facing M1 (**Supplementary Figure 4a-b**), in contrast to the surface-exposed, metal-binding CXXC hallmark-motif detected in classical HMBDs. Functional analysis of mutant forms lacking these cysteines *in vitro* also support that they are unimportant for function (**Figure 2a**). We note that there are P_IB-4_-ATPases with extended N-termini that may harbour such domains^(*16*)^. Furthermore, the sCoaT N-terminus is rich in metal-binding methionine, cysteine, histidine, aspartate and glutamate residues, and this feature is conserved among P_IB-4_-ATPases (**Supplementary Figure 5**). We therefore explored the role of this N-terminal tail through assessment of a sCoaT form lacking the first 33 residues. However, i*n vitro* characterization suggests only minor differences compared to wild-type, indicating that the residues upstream of MA are not essential for catalytic activity (**Figure 2a**). Aggregated, this hints at that no classical HMBD is present, and hence that this level of regulation is absent in P_IB-4_-ATPases, although it cannot be excluded that the N-termini are important *in vivo*.

**Figure 2.**
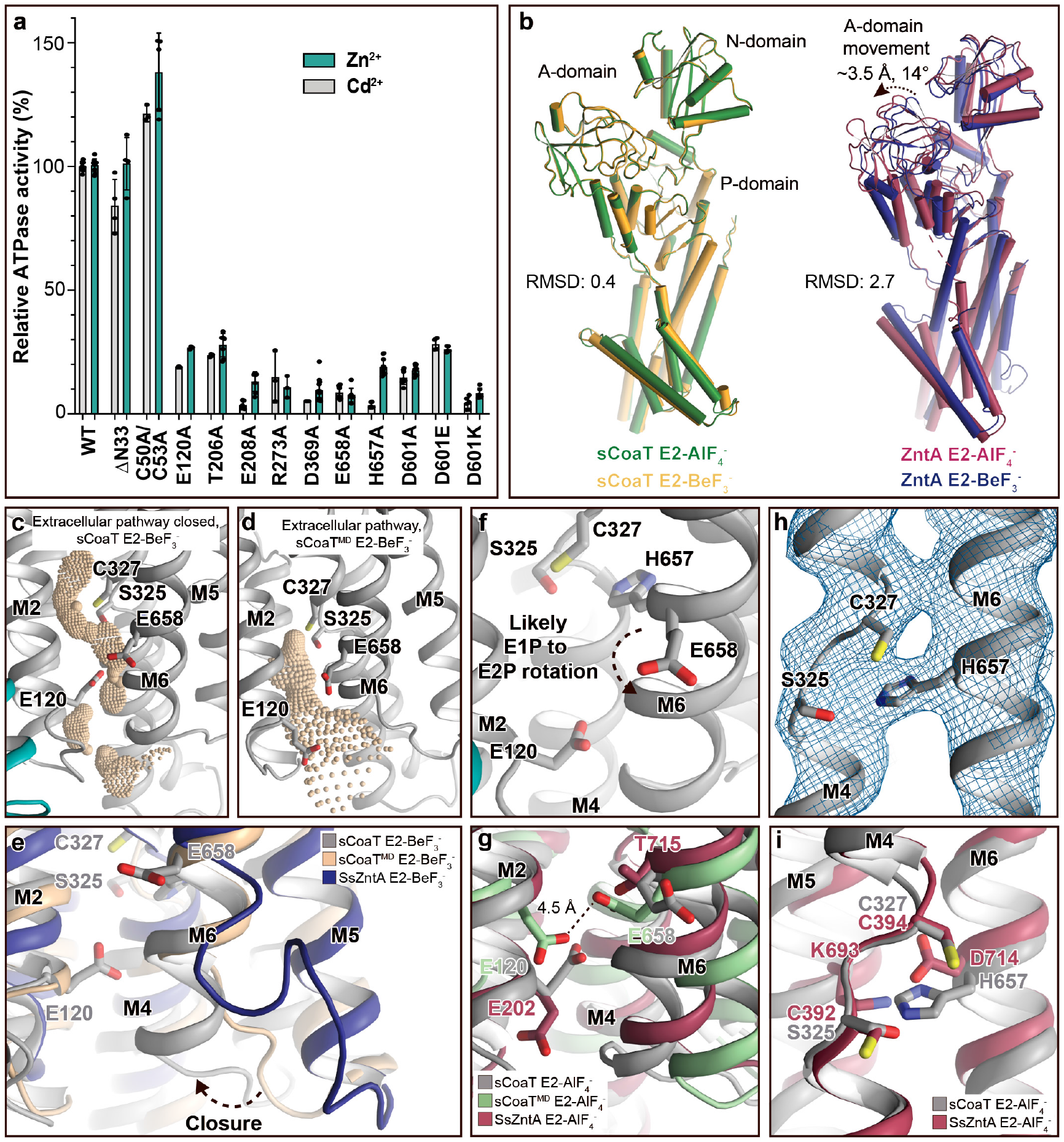
Mechanistic insight into the function of P_IB-4_-ATPases. **a,** Functional ATPase assay in lipid-detergent solution with targeted residues in sequential order. The wild-type (WT) specific activity using the employed experimental conditions in the presence of 50 μM metal is 1.00 ± 0.01 μmol mg^-1^ min^-1^ with Zn^2+^ and 2.80 ± 0.06 μmol mg^-1^ min^-1^ with Cd^2+^, comparable to the activity previously measured for P_IB-4_-ATPases. For biological averages and s.d. see Supplementary Figure 2c. **b**, Comparisons of E2-AlF_4-_ and E2-BeF_3-_ structures of sCoaT and the equivalent of SsZntA (PDB ID of SsZntA structures: 4UMV and 4UMW). All superimpositions were performed based on the overall structures, and the RMSD values for each superimposition are indicated. **c-d**, Identified cavities (wheat) using the software HOLE. **c**, The E2-BeF_3-_ occluded structure lacking connection between the ion-binding site to the outward environment. **d**, The release pathway detected in MD simulations of the E2-BeF_3-_ structure. **e**, The conformational changes that likely allow for closure of the release pathway. Note that the orientation of transmembrane helices M5 and M6 is similar in the E2-BeF_3-_ of SsZntA and following MD simulations of the E2-BeF_3-_ structure of sCoaT. **f-i**, Close-views of ion-binding and -release residues in the M-domain of sCoaT and SsZntA. **f**, The orientation of E658 is incompatible with high-affinity binding, and is likely contributing to ion-release. **g**, Release likely takes place via E658 and E120, residues that are in proximity during the MD simulations. **h**, The sandwiched position between S325 and C327 of H657, including the final 2Fo-Fc electron density (blue). **i**, The position of H657 in sCoaT overlaps with the one of K693 in SsZntA, and both likely serve as in-built counter-ions.

Interestingly, it has been shown that the *in vivo* transport specificity of the sCoaT homolog from *Synechocystis PCC 6803* (CoaT) can be switched from Co^2+^ to Zn^2+^ by exchanging the N-terminal region to that of the Zn^2+^ transporting P_IB-2_ ATPase ZiaA from same organism^(*39*)^. This demonstrates that P_IB-4_-ATPases not only *in vitro* (our data), but also *in vivo* are able to transport Zn^2+^, if the M-domain gain access to the metal. One possible explanation for the change of specificity for the CoaT chimeric construct, is that the N-terminal peptide tail, as also suggested for ATP7B^(*40*)^, prevents ATP hydrolysis through binding to the soluble domains, and this inhibition is then released upon binding of the cognate metal to the N-terminal and/or HMBD. However, it is also possible that the role of the N-terminal region of P_IB-4_-proteins is to impair Zn^2+^ acquisition, an ability that is lost when exchanged with the N-terminal part of ZiaA. From this it is clear that further studies are needed to shed light on the function of the N-terminal region in P_IB_-ATPases, also in P_IB-4_-ATPases.

Associated, this raises questions also on the role of the above-mentioned MB’ platform, which has been proposed to serve as an interaction site for HMBDs in P_IB-1_- and P_IB-2_-ATPases, and for the Cu^+^-ATPases also as a docking site for metal delivering chaperones^(*27, 28, 32, 41*)^. As there are no known zinc/cadmium chaperones for P_IB-4_-ATPases, and because classical HMBDs thus appear to be missing in at least some proteins of the group, the MB’ function may need to be revisited. Alternatively, the N-terminus may have merely been maintained through evolution without conferring functional benefits or disadvantages.

### Structures in transition states of dephosphorylation

The classical view of P-type ATPases is that the E2P state is outward-open and that the following transition state of dephosphorylation, E2.P_i_, is occluded, and that these conformations can be stabilized using the phosphate analogues employed here for structure determination, BeF_3-_ and AlF_4-_, respectively. Furthermore, distinct ion release pathways have been proposed among P_IB_-ATPases^(*27-29, 42*)^, including a narrow exit pathway lined by MA, M2 and M6 that remains open also in the E2.P_i_ state for the P_IB-1_-ATPases. In contrast, a wide opening extending from the location of the bound metal in the M-domain of ion-occluded states to the non-cytoplasmic side has been observed for the P_IB-2_-ATPases, and this group becomes re-occluded with the E2P to E2.P_i_ shift.

Surprisingly however, analysis of the two obtained structures suggests that the anticipated significant domain reorientations are absent in sCoaT (**Figure 2b**), and the models are in contrast rather similar. The compact assembly of the soluble domains and the position of the A-domain near the P-domain, placing the conserved TGE-motif responsible for dephosphorylation towards the phosphorylation site, are typically associated with commencement of dephosphorylation, indicating that the two structures are trapped in E2.P_i_ like transition states (**Supplementary Figure 6a-c)**. This observation differs from the equivalent structures of the other structurally determined P_IB_-ATPases, in which the phosphorylation site of the E2P state (stabilized by BeF_3-_) is shielded from the TGE-loop as also observed for the well-studied sarcoendoplasmic reticulum Ca^2+^-ATPase (SERCA) **(Supplementary Figure 6d)**.

Notably, analogous highly similar BeF_3-_- and AlF_4_-stablized structures have recently also been observed for the Ca^2+^-specific P-type ATPase from *Listeria monocytogenes* (LMCA1)^(*43*)^. It was proposed that LMCA1 pre-organizes for dephosphorylation already in a late E2P state (E2P*, stabilized by BeF_3-_), in accordance with its rapid dephosphorylation. Favoured occlusion and activation of dephosphorylation directly upon ion-release may also be the case for sCoaT, and consequently the E2-BeF_3-_ structure captured here may represents a late (or quasi) E2P state (E2P*).

Comparisons of the sCoaT structures to the equivalent structure of SsZntA (E2.P_i_) revealed a unique arrangement of the A-domain **(Supplementary Figure 7a-b)**. The TGE-loop region superposes well with the corresponding area in SsZntA, but the rest of the A-domain is rotated towards the P-domain - approximately 14° and 3.5 Å **(Supplementary Figure 7b)**. Additionally, we noticed that the A-domain of sCoaT possesses a surface-exposed extension similar to SERCA, but not present in P_IB-1_- and P_IB-2_-ATPases **(Supplementary Figure 7c)**. However, sequence alignments suggest that it is not a conserved feature in the P_IB-4_-group. (**Supplementary Figure 5**). Correspondingly, the M-domains of the two sCoaT structures are overall similar and appear outward-occluded (**Figure 2c)**, as also supported by comparisons with the equivalent structures of SsZntA, again contrasting to the situation observed in P_IB-1_- and P_IB-2_-ATPases.

### Ion-release

Next, to shed light on ion-release, we performed molecular dynamics (MD) simulations using the generated structures embedded in dioleoylphosphatidylcholine (DOPC) lipid bilayers. Strikingly, in the MD simulation of the E2-BeF_3-_ and E2-AlF_4-_ structure of sCoaT, the outward-facing end of M6 changed its orientation (**Figure 2d-e and Supplementary Figure 8a**). This arrangement is similar to the outward-open conformation uncovered in the E2P state of P_IB-2_-ATPases, albeit with a less pronounced opening. Thus, P_IB-2_- and P_IB-4_-ATPases may employ similar exit pathways, lined primarily by M4, M5 and M6, as also substantiated by the overlapping cargo range, and the fact that these two groups release ions in free-form to the extracellular environment, in contrast to their P_IB-1_-counterpart.

The high affinity binding site in P_IB-4_ ATPases has previously been suggested to be formed of residues from the conserved SPC- (starting from S325) and HEGxT (from H657) -motifs of M4 and M6, based on X-ray absorption spectroscopy (XAS) and mutagenesis studies^(*18, 44*)^. An outstanding remaining question is however how the ion then is discharged to the extracellular site? Remarkably, E658 of M6 is pointing away from the ion-binding region around the SPC motif in the sCoaT structures (**Figure 2f and Supplementary Figure 4a**). We anticipate that E658 rotates away from its ion-binding configuration in the E1P to E2P transition, thereby assisting to lower the cargo-affinity to permit release via the M4, M5 and M6 cavity (**Figure 2f**). The conserved E120 of M2 (sometimes replaced with an aspartate in P_IB-4_-ATPases) is located along this exit pathway, and it overlays with the conserved E202 in SsZntA (**Figure 2g**), which has been suggested to serve as a transient metal ligand, stimulating substrate release from the CPC motif of P_IB-2_-ATPases^(*28*)^. We propose a similar role for E120 in sCoaT as further supported by the decreased activity of E120A sCoaT form (**Figure 2a**). In agreement with this notion, E568 of the HEGxT motif and E120 directly interact when the open configuration is obtained in the MD simulations (**Figure 2g and Supplementary Figure 8b**).

### A unique internal counter-ion principle

Many P-type ATPases couple ion- and counter-transport, and hence the reaction cycle cannot be completed without counter-ions. The importance of the counter-transport has been demonstrated in e.g. Ca^2+^/H^+^-(such as SERCA), Na^+^/K^+^- and H^+^/K^+^-ATPases^(*45- 48*)^. In contrast, absence of counter-transport has been proposed for P_IB-2_-ATPases^(*28*)^, H^+^-ATPases^(*49*)^ and P4-ATPases^(*50*)^, which rather exploit a built-in counter-ion Specifically for the P_IB-2_-ATPases, a conserved lysine of M5 (K693 in SsZntA) serves as the counter ion, through interaction with the conserved metal binding aspartate of M6 (D714 in SsZntA) in E2 states. Similarly, P_IB-1_-ATPases are not Cu^+^/H^+^ antiporters, but a likely built-in counter-ion residue is not conserved in the group^(*51*)^. Instead, it is possible that the requirement for counter-ion translocation is prevented by the narrow exit pathway, preventing back-transfer of the released ion and perhaps rendering complete-occlusion unnecessary^(*51*)^. For the P_IB-4_-ATPases, biochemical studies have proposed an ion-binding stoichiometry of one^(*18, 22, 26, 44*)^, however no information is available regarding the presence or absence of counter transport.

In the E2-BeF_3-_ sCoaT structure, we identify a tight configuration of HEGxT-motif H657, being sandwiched between the SPC residues, distinct from the M5 lysine - M6 aspartate interaction observed in P_IB-2_-ATPases (**Figure 2h-i**). Despite the packing issues of the generated crystals, clear electron-density is visible for H657, indicating a rigid conformation (**Figure 2h**). Moreover, activity measurements of an alanine substitution of H657 demonstrate that it is crucial for function (**Figure 2a**). In light of these findings and an earlier report suggesting that a mutation of the equivalent of H657 in MtCtpD leaves the ion affinity unaffected^(*44*)^, we suggest this histidine serves as an internal counter-ion, similarly as for the invariant lysine in SsZntA, perhaps preventing back-transfer of released ions and for charge stabilization, however we cannot exclude that H657 is also part of the high affinity binding site in sCoaT.

The rigid conformation observed for H657 in the E2-BeF_3-_ structure is also observed in the E2-AlF_4-_ structure (**Supplementary Figure 4b**). In contrast, for SsZntA the interaction between K693 and D714 is only detected in the E2.P_i_ state. Thus, the interaction pattern is consistent with the idea that sCoaT pre-organizes for dephosphorylation already in the (late) E2P state, with the associated occlusion and internal counter-ion interaction taking place earlier than for SsZntA.

### A more potent A-domain modulatory site

A conserved K^+^-site, which cross-links between the A- and P-domains in E2 states and thereby allosterically stimulates the E2P to E2 process^(*52, 53*)^, has been suggested to be present also in P_IB_-ATPases^(*52*)^. However, our new E2 structures and available structures of P_IB-1_- and P_IB-2_-ATPases suggest that the A-/P-domain linker is maintained without K^+^ in P_IB_-ATPases, and instead is established directly between R273/D601 in sCoaT as also supported by potassium titration experiments monitoring sCoaT ATPase activity (**Figure 3a-d**). Nevertheless, the point-of-interaction appears critical for P_IB_-ATPases, as functional characterization of R273A, D601A and D601K result in a marked reduction of turn-over (**Figure 2a**). This differs from similar mutations of classical P-type ATPases, where only minor effects are observed^(*52, 53*)^. Furthermore, substitution of D601 with glutamate suggests that even the A-/P-domain distance is critical (**Figure 2a**). It is possible that P_IB_-ATPases are more reliant on this particularly tight, ion-independent stabilization, as the A-M1/A-domain linker is absent, and because many other P-type ATPases also have a complementary A-/P-domain interactions. Thus, our data indicate that this regulation is a general feature of many P-type ATPase classes, yet featuring unique properties for P_IB_-ATPases.

**Figure 3.**
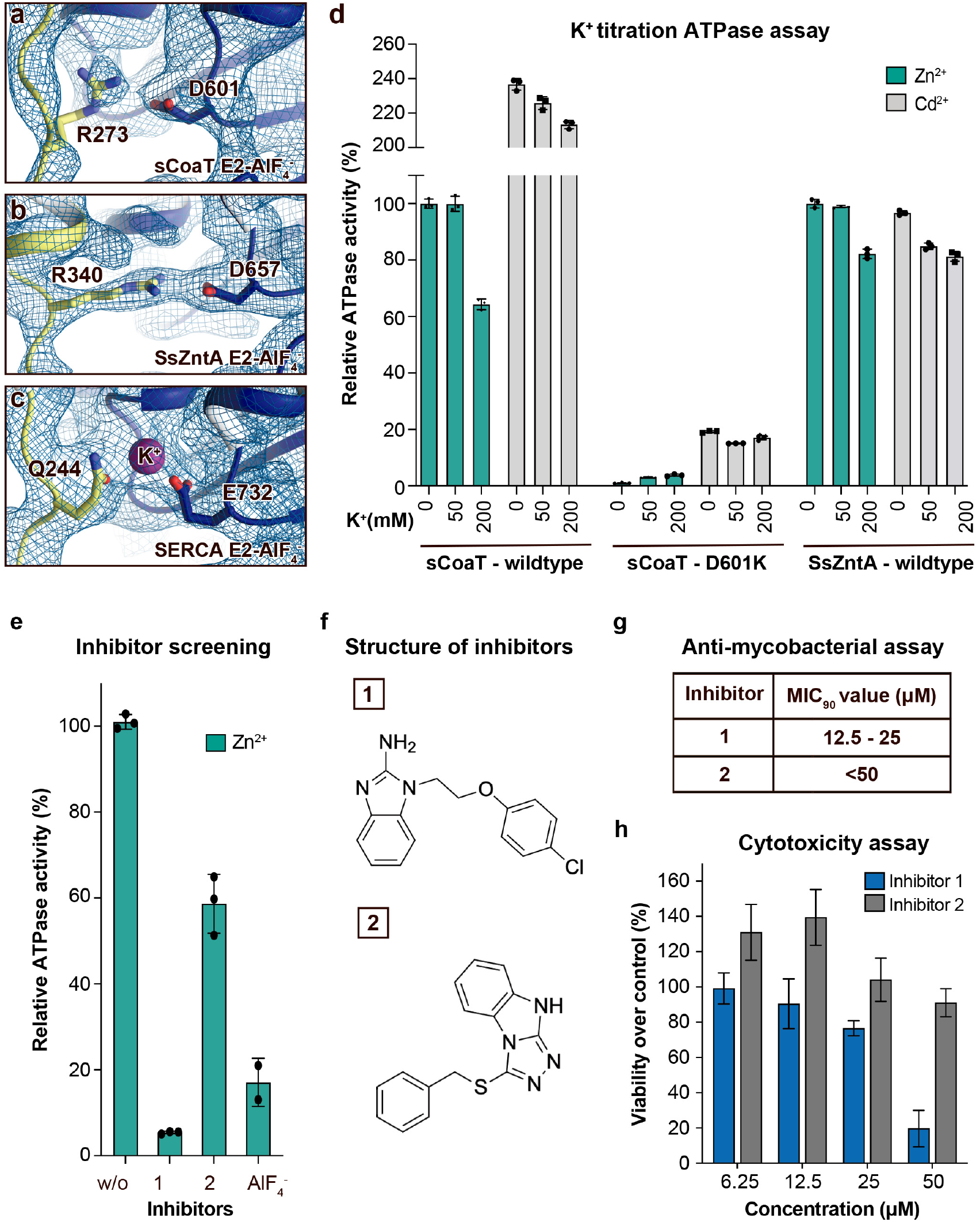
Regulation and inhibition. **a-c,** Close-views of the regulatory point of interaction between the A- and P-domains in the E2-AlF_4-_ structures of sCoaT, SsZntA and SERCA (PDB IDs 4UMW and 1XP5) with the corresponding 2Fo-Fc electron density shown at σ=1.0 (blue mesh). **a**, sCoaT (colored as in Figure 1) with interaction between D601 and R273. **b**, SsZntA (shown as panel a) with interaction between D657 and R340. **b**, SERCA (shown as in panel a) with bound K^+^ (purple) between E732 and Q244. **d**, Functional ATPase assay in lipid-detergent solution of sCoaT (wild-type and D601K forms) as well as SsZntA (wild-type), using protein samples purified in the absence of K^+^ and Na^+^ (see Methods). The mean + s.d. of technical replicates is shown (n=3). KCl leaves the function of sCoat and SsZntA essentially unaffected in the presence of Zn^2+^ (cyan) or Cd^2+^ (gray). The equivalent form of sCoaT D601K has previously been exploited to demonstrate K^+^-dependence in the Na,K-ATPase^(*53*)^. Collectively, these data suggest that the P-/A-domain site regulation is K^+^-independent in P_IB_-ATPases, in contrast to classical P-type ATPases. **e-h** Evaluation of the effect on selected identified novel inhibitors on activity of protein, as well as survival of mycobacteria and primary human macrophages. **e**, Effect of two inhibitors (300 μM) on the activity of sCoaT assessed in lipid-detergent solution in the presence of Zn^2+^. For comparison, the commonly used P-type ATPase inhibitor AlF_4-_ (500 μM) is included. **f**, The structure of inhibitor 1 and 2. **g**, The minimal inhibitory concentration to kill 90 % (MIC_90_) of mycobacteria for inhibitor 1 and 2. The mean MIC_90_ value for inhibitor 1 is 18.75 µM, while for inhibitor 2 it is over 50 µM. The values are based on four separate experiments. **h**, The cytotoxic effect of different concentrations of inhibitor 1 and 2 on primary human macrophages (ATP assay). The standard error of mean (SEM) of 9 replicates is shown (n=9).

### New metal-transport blockers

P_IB-2_- and P_IB-4_-ATPases serve as virulence factors and are critical for the disease caused by many microbial pathogens, as underscored by the frequent presence of several redundant genes^(*54-57*)^. In this light and because these P-type ATPases are missing in humans, they represent putative targets for novel antibiotics. The shared mechanistic principles identified here suggest that compounds can be identified that inhibit both P_IB_-groups, for example directed against the common release pathway, thereby increasing efficacy. Indeed, screening of a 20,000-substance library using a complementary *in vitro* assay, uncover several leads that abrogate function of sCoaT and SsZntA (**Figure 3e-f**, data only shown for sCoaT). Furthermore, initial tests of two of these suggest they have a potent effect against mycobacteria, which previously have been shown to be P_IB-4_-dependent for infection^(*44*)^; 90 % of the mycobacteria were killed at mean concentrations of 18.75 and above 50 µM, respectively, using either of these two separate molecules (**Figure 3g**). In contrast, investigation of cytotoxic effects on primary human macrophages at concentrations up to 25 µM demonstrated considerably less impact on cell survival for both blockers (**Figure 3h**). Evidently downstream in-depth studies, ranging from investigations of the target specificity, the detailed effect on human cells as well as antibiotic potency in human, are required to fully understand the value of these putative P_IB-2_- and P_IB-4_-inhibitors. Nevertheless, the substances outlined here represent promising leads for drug-discovery efforts or to aid the development of tools to manipulate heavy metal accumulation in plants to prevent accumulation or for enrichment.

## Conclusion

Collectively, the first structure of a P_IB-4_-type ATPase reveals the topology of P_IB-4_-ATPases, displaying an eight helix M-domain configuration, and likely no HMBDs, at least in members without extended N-termini. Major findings include the observation of an ion-release-pathway similar as in the related P_IB-2_-ATPases, a previously not observed counter-ion principle for P-type ATPases, and a unique potassium-independent regulation of the P_IB_-transport cycle (**Figure 4**). Thus, our results significantly increase the understanding of heavy metal homeostasis in cells. The novel identified putative inhibitors and the partially overlapping mechanistic principles of P_IB-2_- and P_IB-4_-ATPases also open up a novel avenue for development of compounds accessible from outside the cell against these P_IB_-groups, to combat global threats such as multi-drug resistance and/or tuberculosis or for biotechnological purposes.

**Figure 4.**
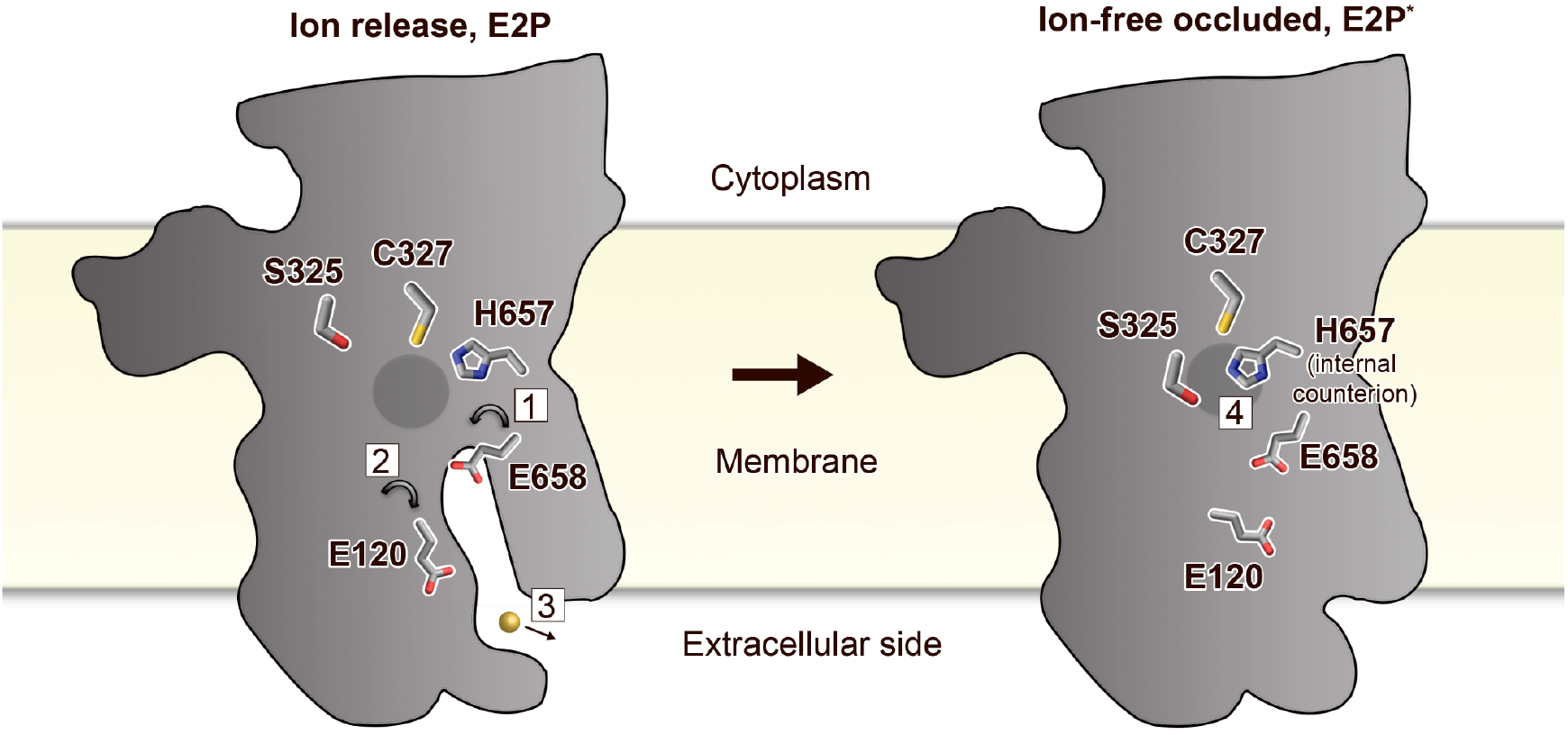
Putative ion-release and re-occlusion mechanism of P_IB-4_-ATPases. Schematic model illustrating the transmembrane domain (the soluble domains have been removed for clarity) of two separate states, an E2P and an occluded E2P* conformation as the determined structure (E2-BeF_3-_), respectively. Zinc or cadmium release from the high affinity binding site in the M-domain is likely permitted through re-orientation of E658 (1) in the E1P to E2P transition, thereby lowering the affinity for the occluded ion. E120 serves as a transient linker between the high-affinity binding site and the outward environment (2). Following ion-release (3) H657 shifts to a sandwiched position between S325 and C327 (4), acting as a built-in counter ion, preventing back-transfer of the released ion, and allowing completion of the reaction cycle.

## Materials and methods

### Overproduction and purification of sCoaT

Forms of the 72 kDa sCoaT from *Sulfitobacter* sp. NAS14-1 (UniProt ID A3T2G5) were transformed into *E. coli* (C41 strain) cells. The cells were cultured in LB medium at 37 °C with shaking at 175 rpm in baffled flasks until the optical density (600 nm) reached 0.6-1, cooled to 18 °C, and then induced with 1 mM IPTG for 16h. Harvested cells were resuspended in buffer A (1 g cells per 5 mL buffer) containing 20 mM Tris-HCl, pH=7.6, 200 mM KCl, 20 % (v/v) glycerol and frozen at -80 °C until further use. Cells were disrupted by two runs in a high-pressure homogenizer (Constant System) at 25,000 psi following addition of 5 mM of fresh β-mercaptoethanol (BME), 5 mM MgCl_2_, 1 mM phenylmethanesulphonyl fluoride, 2 μg/mL DNase I and Roche protease inhibitor cocktail (1 tablet for 6 L cells). The sample was kept at 4 °C throughout the purification. Cellular debris was pelleted via centrifugation at 20,000 g for 20 minutes. Membranes were isolated by ultracentrifugation for 3 h at 185,500 g, and resuspended in 10 mL buffer B (20 mM Tris-HCl, pH=7.6, 200 mM KCl, 1 mM MgCl_2_, 5 mM BME and 20 % (v/v) glycerol) per g membranes and frozen at -80 °C until further use. The protein concentration in the membranes was estimated using the Bradford assay^(*58*)^. Proteins were solubilized through supplementation of 1 % (w/v) final concentration n-dodecyl-β-D-maltopyranoside (DDM) and 3 mg/mL final total protein concentration in Buffer B with gentle stirring for 2 h. Un-solubilized material was removed by ultracentrifugation for 1 h at 185,500 g. The supernatant was supplemented with imidazole to a final concentration of 30 mM and solid KCl (500 mM final concentration), filtered (0.22 mm) and then applied to 5 mL HiTrap Chelating HP columns (GE Healthcare, protein from 6 L cells per column) charged with Ni^2+^ and equilibrated with 4 column volumes of buffer C (20 mM Tris-HCl, pH=7.6, 200 mM KCl, 1 mM MgCl_2_, 5 mM BME, 150 mg/mL octaethylene glycol monododecyl ether (C_12_E_8_) and 20 % (v/v) glycerol). Proteins were eluted using a gradient, ending with buffer C containing 500 mM imidazole. Eluted protein was assessed using SDS–PAGE, and the fractions containing sCoaT concentrated to approximately 20 mg/mL using VivaSpin concentrators (MWCO=50 kDa). 10 mg concentrated protein was subjected to size-exclusion chromatography using a Superose 6 gel-filtration column (GE-Healthcare), pre-equilibrated with 50 mL buffer E (20 mM Tris-HCl, pH=7.6, 80 mM KCl, 1 mM MgCl_2_, 5 mM BME, 150 mg/mL C_12_E_8_ and 20 % (v/v) glycerol). Fractions containing purified sCoaT were pooled, and concentrated to approximately 10 mg/mL, flash frozen in liquid nitrogen, and stored at -80 °C until further use. For the experiments to assess K^+^-dependence, the buffer E was replaced with 20 mM Tris-HCl, pH=7.5, 1 mM MgCl_2_, 5 mM BME, 0.15 mg/mL C_12_E_8_ and 20 % (v/v) glycerol.

### Crystallization

10 mg/mL sCoaT was supplemented with 3 mg/mL (final concentration) DOPC and 6 mg/mL (final concentration) C_12_E_8_, incubated at 4 °C and stirring for 16-48 h (modified HiLiDe method^(*36*)^). Aggregates and insoluble DOPC were then removed by ultracentrifugation at 50,000 g for 10 minutes. 2 mM AlCl_3_ or BeSO_4_, 10 mM NaF and 2 mM EGTA (final concentrations) were supplemented and incubated on ice for 30 minutes. Crystals were grown using the hanging drop vapor diffusion method at 19 °C. E2-AlF_4-_ crystals were grown with a reservoir solution containing 200 mM MgCl_2_, 14 % (v/v) PEG1500, 10 mM tris(2-carboxyethyl)phosphine, 10 % (v/v) glycerol, 3 % 2-Methyl-2,4-pentanediol and 100 mM sodium acetate, pH=5.0. The E2-BeF_3-_ crystals were grown with a reservoir solution containing 200 mM magnesium formate, 14 % (v/v) PEG5000, 100 mM sodium acetate, pH=4.0, and 0.5 % (v/v) 2-propanol was added as an additive. Crystals were fished using litholoops (Molecular Dimensions), flash-cooled in liquid nitrogen, and tested at synchrotron sources. Complete final data sets were collected at the Swiss Light Source, the Paul Scherrer Institute, Villigen, beam line X06SA.

### Structure determination and refinement

Collected data were processed and scaled with XDS (**Supplementary Table 1)**. For the E2-AlF_4-_ structure, initial phases were obtained by the molecular replacement (MR) method using software PHASER^(*59*)^ of the Phenix package^(*60*)^, and using the AlF_4-_-stabilized structure of SsZntA (PDB ID: 4UMW) as a search model. The E2-BeF_3-_ structure was solved using the generated E2-AlF_4-_ structure as a MR model. Both crystal forms display poor crystal packing between the membrane domains (**Supplementary Figure 3**), deteriorating the quality of the electron density maps in these regions (**Supplementary Figure 4**). In this light, model building of the membrane domains were executed with particular prudence, taking into consideration the connectivity to the well-resolved soluble domains, distinct structural features as well as sequence and structure conservation patterns. Examples of such include the conserved GG motif that forms the kink in MB helix, which is clearly identified also at low resolution, the SPC motif that twists the M4 helix and the conserved and functionally important well-resolved residue H657 that assisted assigning nearby residues.

Initial manual model building was performed primarily using COOT^(*61*)^. ISOLDE^(*62*)^ in ChimeraX^(*63*)^ was employed for model building and analysis, and was critical for obtainment of the final models with reasonable chemical restraints and low clash score. In particular, ISOLDE’s interactive register shifting tool was instrumental in determining the register of the most weakly resolved TM helices. Secondary structure restraints were applied in some flexible regions, also taking into consideration homology to sCoaT and other models.

During final refinements with phenix.refine^(*64*)^ the geometry was restrained in torsion space to ISOLDE’s output. Molprobity was exploited for structure validation^(*65*)^. The final models are lacking the first 40 residues only, which is shorter than a classical MBD of 67 amino acids. All structural figures were generated using Pymol^(*66*)^. Statistics for the final models were 96.70, 3.30, 0.20 and 0,74 for E2-BeF_3-_ and 93.24, 6.13, 0.63 and 8.31 for E2-AlF_4-_ in Ramachandran favored and allowed regions, and for rotamer outliers and clash-score, respectively.

### Activity assay

sCoaT forms were functionally characterized using the Baginski method to assess the amount of released inorganic phosphate^(*67*)^. Briefly, 0.5 μg of purified sCoaT mixed with reaction buffer containing 40 mM MOPS-KOH, pH=6.8, 5 mM KCl, 5 mM MgCl_2_, 150 mM NaCl, 0.3 mg/mL C_12_E_8_, 0.12 mg/mL soybean lipid, 5 mM NaN_3_ and 0.25 mM Na_2_MoO_4_ in a total volume of 50 uL. For metal stimulation assays, different heavy metal ions or EGTA was supplemented the reaction buffer to a final concentration of 50 µM. For inhibitor screening (see how inhibitors were identified below), different concentrations of inhibitors were added to the reaction buffer containing 50 μM ZnCl_2_. The samples were then incubated at 37 °C with 500 rpm shaking for 5 minutes, and then supplemented with 5 mM ATP (final concentration) to start the reaction, and incubated at 37 °C with 1000 rpm shaking for 10 mins. 50 μL freshly prepared stop solution containing 2.5 % (w/v) ascorbic acid, 0.4 M (v/v) HCl and 1 % SDS was then supplemented to stop the reaction and start colour development. 75 μL color solution (2 % (w/v) arsenite, 2 % (v/v) acetic acid and 3.5 % (w/v) sodium citrate) was added to the mixture following 10 minutes incubation at room temperature. Absorbance was measured at 860 nm after another 30 minutes incubation at room temperature. For the experiments to assess K^+^-dependence, the reaction buffer was replaced with 40 mM Tris-HCl, pH=7.5, 5 mM MgCl_2_, 3.0 mg/mL C_12_E_8_ and 1.2 mg/mL soybean lipid in a total volume of 50 uL.

### Inhibitor screening

The inhibitor screening experiments were initially carried out on the zinc transporting P_IB-2_-type ATPase ZntA from *Shigella sonnei* (SsZntA). SsZntA was produced and purified as described previously^(*28*)^ and the inhibitory effect of approximately 20000 compounds was assessed by the Chemical biology Consortium Sweden (CBCS). Briefly, the ATPase activity of 0.7 µM highly pure protein was measured in the presence of 50 µM inhibitor through the release of inorganic phosphate (P_i_) by the Baginski assay^(*67*)^ in a total volume of 200 nL as reported earlier^(*28*)^. The inorganic phosphate was detected with Malachite Green reagent (0.005 % Carbinol hydrochloride, 1.7 % sulfuric acid, 0.14 % ammonium molybdate, 0.025 % Triton-X) at an absorbance of 620 nm.

### Minimum inhibitory concentration (*MIC*_*90*_)

*Mycobacterium bovis* bacillus Calmette-Guerin (BCG) Montreal containing the pSMT1-*luxAB* plasmid was prepared as previously described^(*68*)^. Briefly, the mycobacteria were grown in Middlebrook 7H9 broth, supplemented with 10% ADC enrichment (Middlebrook Albumin Dextrose Catalase Supplement, Becton Dickinson, Oxford, UK) and hygromycin (50 mg/l; Roche, Lewes, UK), the culture was washed twice with sterile PBS, and re-suspended in broth and then dispensed into vials. Glycerol was added to a final concentration of 25% and the vials were frozen at −80°C. Prior to each experiment, a vial was defrosted, added to 9 mL of 7H9/ADC/hygromycin medium, and incubated with shaking for 72 h at 37°C. Mycobacteria were then centrifuged for 10 minutes at 3000×g, washed twice with PBS, and re-suspended in 10 ml of PBS. Resazurin microtiter assay (REMA) was used to determine the minimum inhibitory concentration (MIC_90_) for the inhibitors against the mycobacterial strain. The inhibitors (10 µL) were added to bacterial suspensions (90 µL) on a 96-well plate at a concentration range between 0.4-50.0 µM. MIC was determined by the color change using resazurin (1:10 v/v, PrestoBlue Cell viability reagent, Thermo Scientific). MIC was determined after one week by adding 10 µL resazurin followed by incubation overnight, corresponding to 90% inhibition.

### Human cytotoxicity assays

Human venous blood mononuclear cells were obtained from healthy volunteers using a Lymphoprep density gradient (Axis-Shield, Oslo, Norway) according to the manufacturer’s instructions. To obtain pure monocytes, CD14 micro beads were applied to the cell suspension, washed and passed through a LS-column according to manufacturer’s description (130-050-201, 130-042-401, Miltenyi Biotec, USA). The monocytes were counted (Sysmex), diluted in RPMI 1640 supplemented with 5% FCS, NEAA, 1 mM Sodium Pyruvate, 0.1 mg/mL Gentamicin (11140-035, 111360-039, 15710-49, Gibco, Life Technologies) and 50 ng/mL GM-CSF (215-GM, R&D systems) and seeded in 96-well plates (10^5^/well) for a week to differentiate into macrophages. Infection experiments were performed in RPMI 1640 without Gentamicin. The medium was replaced with fresh medium containing 6.3, 12.5, 25 or 50 μM inhibitor or DMSO and incubated 24 h in 5% CO_2_ atmosphere. For cytotoxicity measurement, 10 μL 3-(4,5-dimethylthiazol-2-yl)-2,5 diphenyltetrazolium bromide (MTT) solution (Sigma) was added to each well according to manufacturer’s instructions and analysed in a spectrophotometer at 580 nm. NZX cytotoxicity was further examined by ATPlite™ assays. Primary macrophages were treated with 6.3, 12.5, 25 or 50 μM inhibitor or DMSO (Sigma) for 24 hours. Cell viability was assessed with cellular ATP levels using ATPlite™ kit (6016943, Perkin Elmer) compared to untreated controls, according to the manufacturer’s instructions.

### MD simulation

The two crystal structures, E2-AlF_4-_ and E2-BeF_3-_, were inserted into a DOPC (1,2-dioleoyl-sn-glycero-3-phosphocholine) membrane patch using the CHARMM-GUI membrane builder^(*69*)^. The membrane positions were predicted by the Orientations of Proteins in Membranes (OPM) server^(*70*)^. During the simulation equilibration phase, position restraints were gradually released from the water and lipids for a total of 30 ns followed by 500 ns non-restrained production runs. Each protein state was simulated in independent repeat simulations starting from a different set of initial velocities, adding up to a sampling total of 500 ns x 4. A Nose-Hoover temperature coupling^(*71*)^ was applied using a reference temperature of 310 K. A Parrinello-Rahman pressure coupling^(*72*)^ was applied with a reference pressure of 1 bar and compressibility of 4.5e-5 bar^-1^ in a semi-isotropic environment. The TIP3P water model was used and the system contained 0.15 M NaCl. The E2-AlF_4-_ system was composed of 256 lipids and 29429 water molecules while E2-BeF_3-_ system was composed of 254 lipids and 30535 water molecules. The systems were equilibrated and simulated using the GROMACS-2021 simulation package^(*73*)^ and CHARMM36 all-atom force fields^(*74*)^ for the protein and lipids. The HOLE analyses were based on representative averages from the final 250 ns of the trajectories.

## Acknowledgements

CG is currently paid by The BRIDGE - Translational Excellence Programme at University of Copenhagen funded by the Novo Nordisk Foundation (NNF18SA0034956). The PhD studies of CG were partly financed by “The memorial foundation of manufacturer Vilhelm Pedersen and wife – and the Aarhus Wilson consortium”. QH was supported by China Scholarship Council. DRM was funded by Carl Tryggers foundation (CTS 17:22), MA was funded by a Swedish Research Council Starting Grant (2016-03610). The computations were performed on resources provided by the Swedish National Infrastructure for Computing (SNIC) through the High-Performance Computing Center North (HPC2N) under project SNIC 2018/2-32 and SNIC 2019/2-29. PG is supported by the following Foundations: Lundbeck, Knut and Alice Wallenberg, Carlsberg, Novo-Nordisk, Brødrene Hartmann, Agnes og Poul Friis, Augustinus, Crafoord as well as The Per-Eric and Ulla Schyberg. Funding is also obtained from The Independent Research Fund Denmark, the Swedish Research Council and through a Michaelsen scholarship. GM is supported by the Robert A. Welch Foundation (AT-1935-20170325; AT-2073-20210327), the National Institute of General Medical Sciences of the National Institutes of Health (R35GM128704) and the National Science Foundation (CHE 2045984). GG is funded by the Swedish Heart-Lung Foundation (20200378), Alfred Österlunds Foundation, Royal Physiographic Society of Lund. We are grateful for assistance with crystal screening at PETRA III at DESY, a member of the Helmholtz Association (HGF), beamline P13, and crystal screening and data collection at the Swiss Light Source, the Paul Scherrer Institute, Villigen, beam line X06SA. Access to synchrotron sources was supported by the Danscatt program of the Danish Council of Independent Research. We acknowledge the Chemical Biology Consortium Sweden (CBCS) at Umeå University that performed the ligand screening.

## Author Contributions

CG and PG contributed to the identification of the scientific problem, and CG initiated the project. CG, QH and NS performed cloning, overproduction, purification and activity measurements. CG and QH accomplished the crystals. CG, QH and PG collected diffraction data, KW solved the structures, CG, QH and KW refined the structures, TC finalized the structures using ISOLDE. AD produced protein for ligand screening and JE was responsible for the screening of the ligands at the Chemical Biology Consortium Sweden. EL performed the initial selection of ligands from the high throughput screening. EL screened the ligands against SsZntA, and QH against sCoaT. DRM conducted MD simulations, supervised by MA. KUR and DIH performed the cellular studies of the inhibitors, supervised by GG. CG prepared the figures assisted by QH and EL. CG, QH, EL and PG contributed to the data analysis and interpretation. CG and PG wrote the first draft. All authors commented on the manuscript. CG and PG supervised the project.

## Data Information

Atomic coordinates and structure factors for the sCoaT AlF_4-_- and BeF_3-_-stabilized crystal structures have been deposited at the Protein Data Bank (PDB) under accession codes XXXX and XXXX. The authors declare no competing financial interests. Correspondence and requests for materials should be addressed to P.G.

## Supplementary information

**Supplementary Table 1.** Data collection and refinement statistics. Statistics for the highest resolution shell are shown in parentheses.

|  | E2-BeF<sub>3</sub><sup>-</sup> | E2-AlF<sub>4</sub><sup>-</sup> |
| --- | --- | --- |
| **Data collection** |  |  |
| Wavelength (Å) | 1.0 | 1.0 |
| Space group | P 21 21 2 | P 21 21 2 |
| Cell dimensions |  |  |
| *a*, *b*, *c* (Å) | 89.0 94.5 128.8 | 89.6 93.7 128.3 |
| $\alpha$, $\beta$, $\gamma$ (°) | 90 90 90 | 90 90 90 |
| Resolution (Å) | 47.3-3.1 | 45.6-3.3 |
|  | (3.22-3.11) | (3.37-3.25) |
| *R*<sub>merge</sub>(%) | 11.4 (276.3) | 15.5 (246) |
| *I* / $\sigma I$ | 17.8 (1.12) | 8.5 (0.98) |
| CC<sub>1/2</sub> | 1 (0.475) | 0.99 (0.37) |
| Completeness (%) | 97.3 (99.8) | 99.2 (99.9) |
| Redundancy | 13.3 (13.8) | 6.1 (6.6) |
| **Refinement** |  |  |
| Resolution (Å) | 47.3-3.1 | 45.6-3.3 |
|  | (3.22-3.11) | (3.37-3.25) |
| No. reflections | 19643 (1963) | 17466 (1714) |
| *R*<sub>work</sub> / *R*<sub>free</sub> (%) | 24.4/26.8 | 21.8/25.5 |
| *No. of atoms* |  |  |
| Protein | 4695 | 4695 |
| Ligand/ion | 5 | 6 |
| Water | 10 | 0 |
| *Average B-factors* |  |  |
| Protein | 135.91 | 152.54 |
| Ligand/ion | 84.15 | 86.47 |
| Solvent | 79.62 |  |
| *R.m.s. deviations* |  |  |
| Bond lengths (Å) | 0.004 | 0.003 |
| Bond angles (°) | 0.77 | 0.83 |
| *Ramachandran statistics* |  |  |
| Favored (%) | 97.8 | 96.9 |
| Allowed (%) | 2.2 | 3.1 |
| Outliers (%) | 0.0 | 0.0 |
| Clashscore | 1.05 | 7.89 |
| MolProbity score | 0.85 | 1.62 |

**Supplementary Figure 1.**
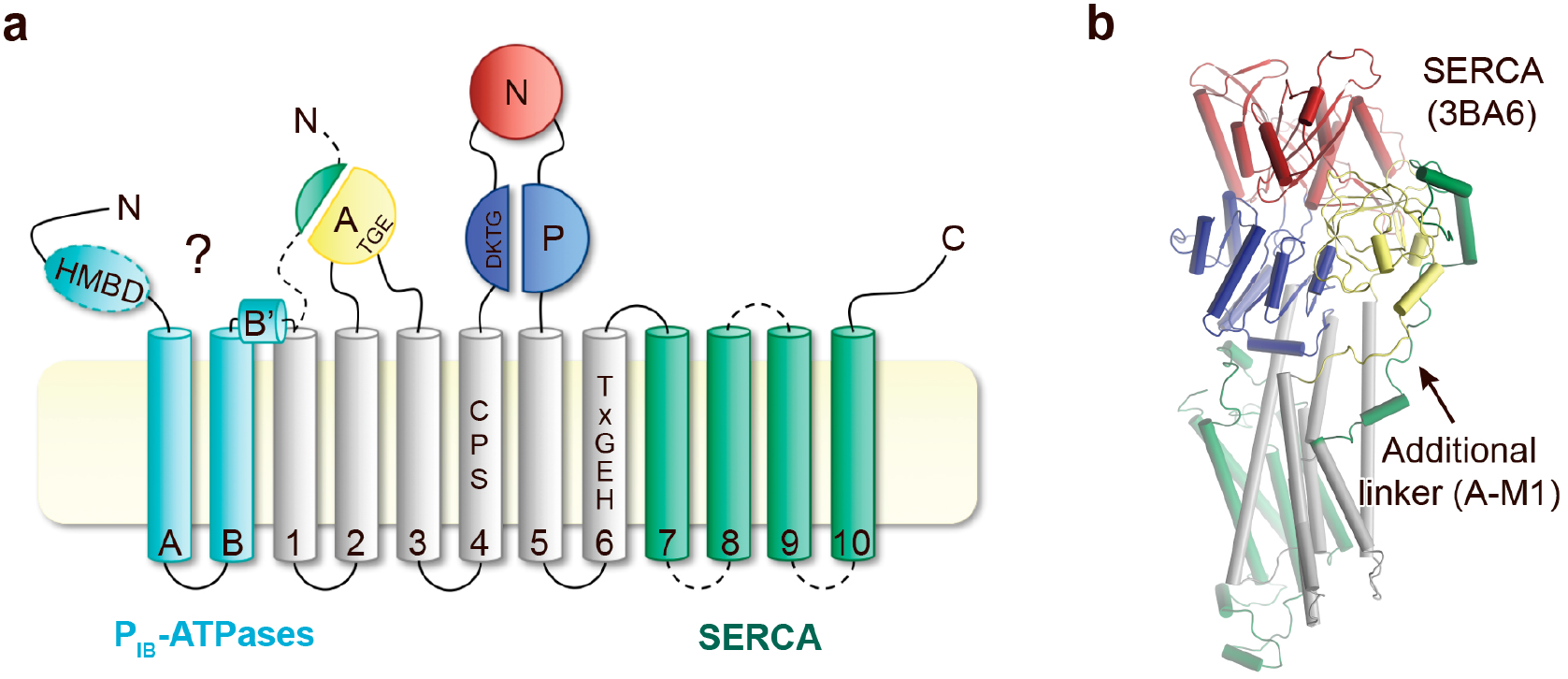
Topology comparison. Topological differences between P_IB_-ATPases and classical P-type ATPases such as sCoaT and SERCA, respectively. **a**, Schematic topology of P-type ATPases showing features unique to P_IB_-ATPases (the so-called heavy metal binding domain, HMBD, and transmembrane helices MA-MB in cyan) and SERCA (an extended A-domain, an additional A-domain linker and M7– M10 transmembrane helices in green). The present work discloses the presence of helices MA, MB, MB’ and that the core of P_IB-4_-ATPases is devoid of classical HMBD, matters that were elusive previously. Key residues in the M-domain for P_IB-4_-ATPases are highlighted. **b**) The structure of SERCA (PDB ID 3BA6), colored as the schematic topology highlighting the additional linker to the A-domain.

**Supplementary Figure 2.**
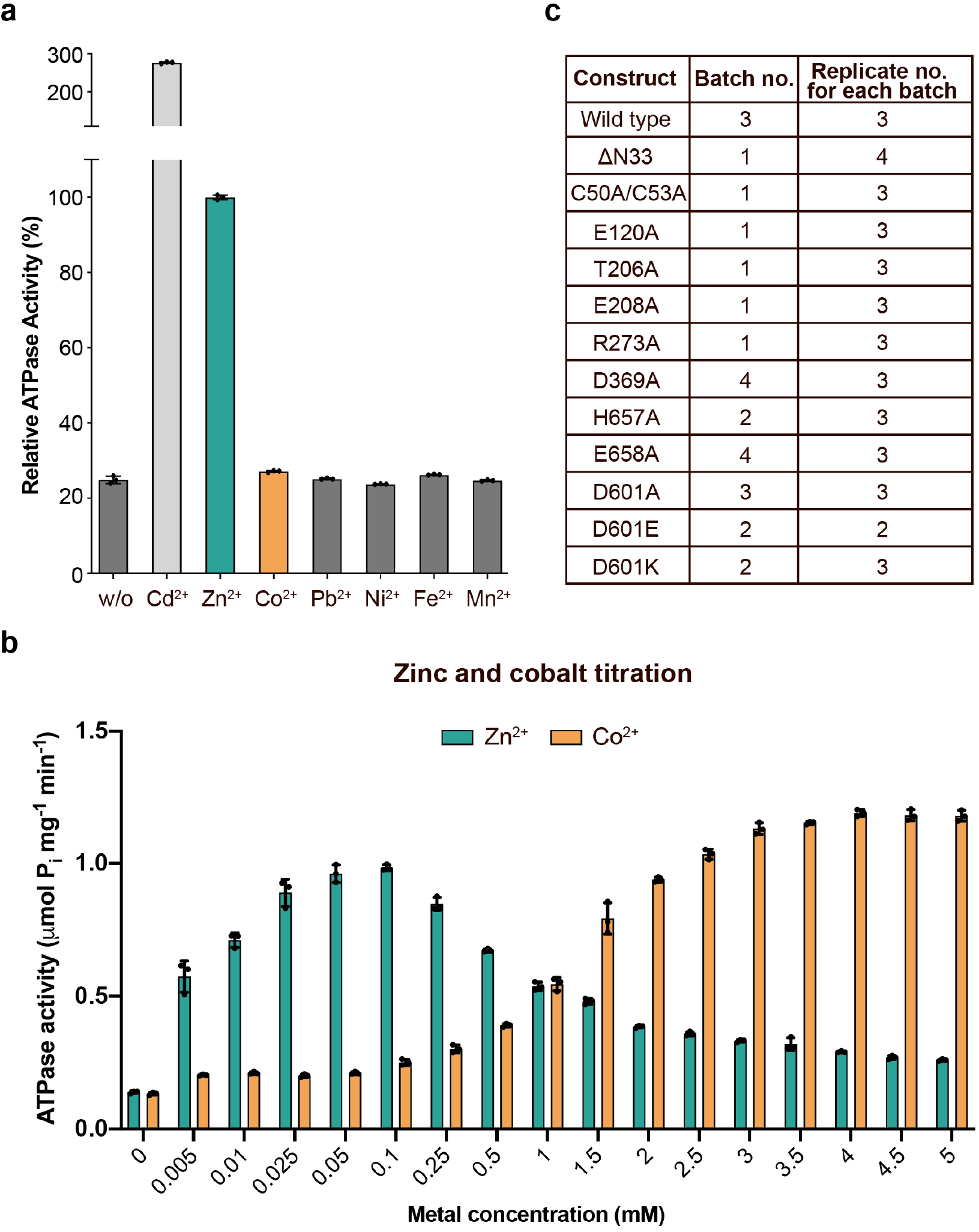
Metal selectivity screening and reproducibility. a, Screen of different metals, tested at 50 μM each. There is clear activity with zinc (1.00 ± 0.01 μmol mg^-1^ min^-1^) and cadmium (2.81 ± 0.05 μmol mg^-1^ min^-1^), while only low ATPase activity (about 5 % of the wild-type, corrected for the background observed with no metal added) was detected with cobalt. **b**, Titration of zinc and cobalt. Cobalt-induced ATPase activity predominates above 1 mM, while at lower concentrations zinc stimulates activity at a faster rate. **c**, Biological and technical replicates exploited to generate the error bars in Figure 2a and Supplementary Figure 6e.

**Supplementary Figure 3.**
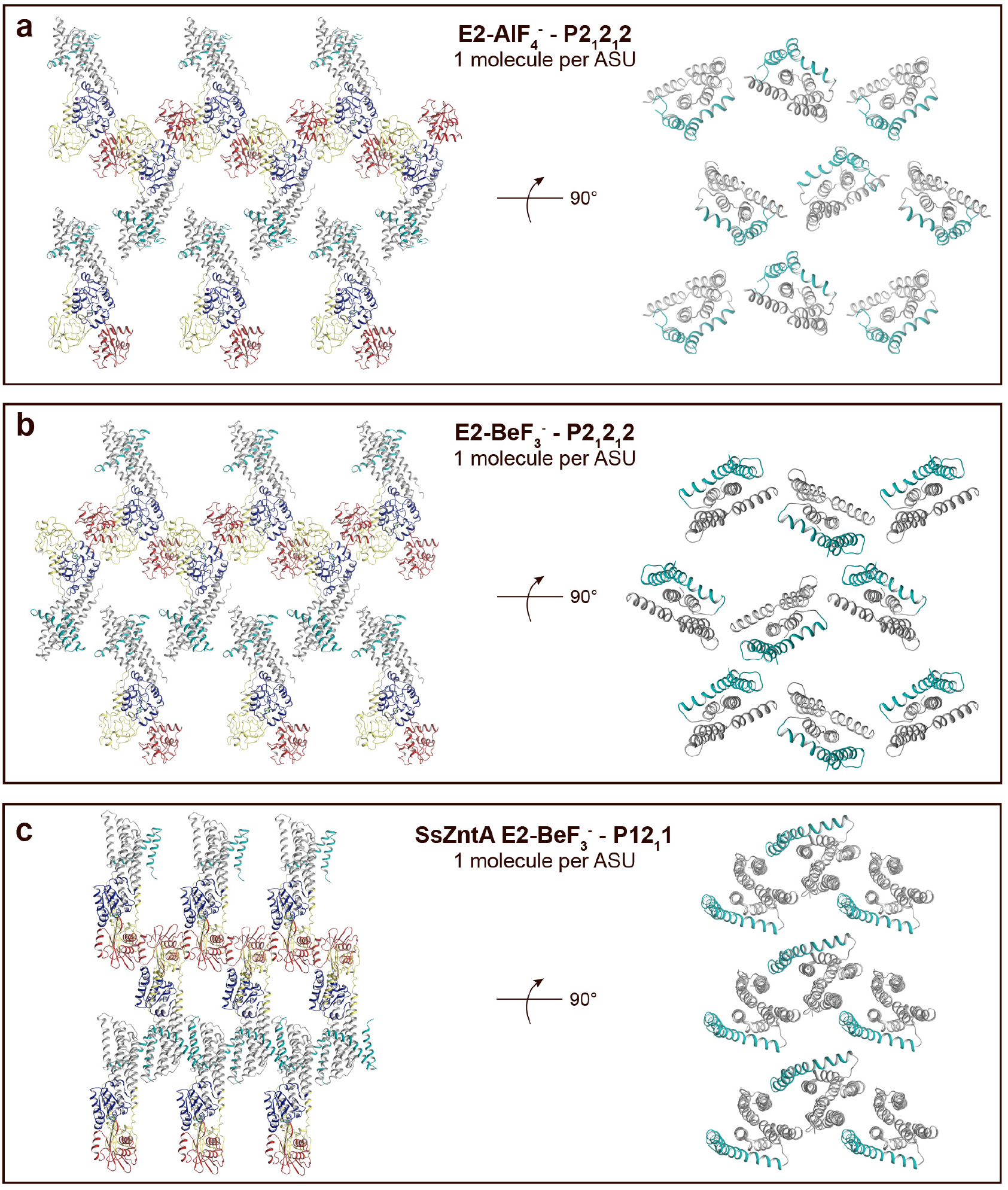
Crystal packing of sCoaT E2-AlF_4-_ compared to the E2-BeF_3-_ crystal form of ZntA from *Shigella sonnei* (SsZntA, PDB ID: 4UMV). The domains are coloured as in Figure 1b. **a**, sCoaT E2-AlF_4-_ (P2_1_2_1_2, with 1 molecule per asymmetric unit). Left: View of the membrane layer. Right: 90° rotation view (compared to panel to the left) showing only the transmembrane domains. Equivalent views of the sCoaT E2-BeF_3-_ (P2_1_2_1_2) (**b**) and the SsZntA E2-BeF_3-_ (P12_1_1) (**c**) crystal forms. Note the loose packing of the sCoaT crystals compared to that of SsZntA E2-BeF_3_^-^.

**Supplementary Figure 4.**
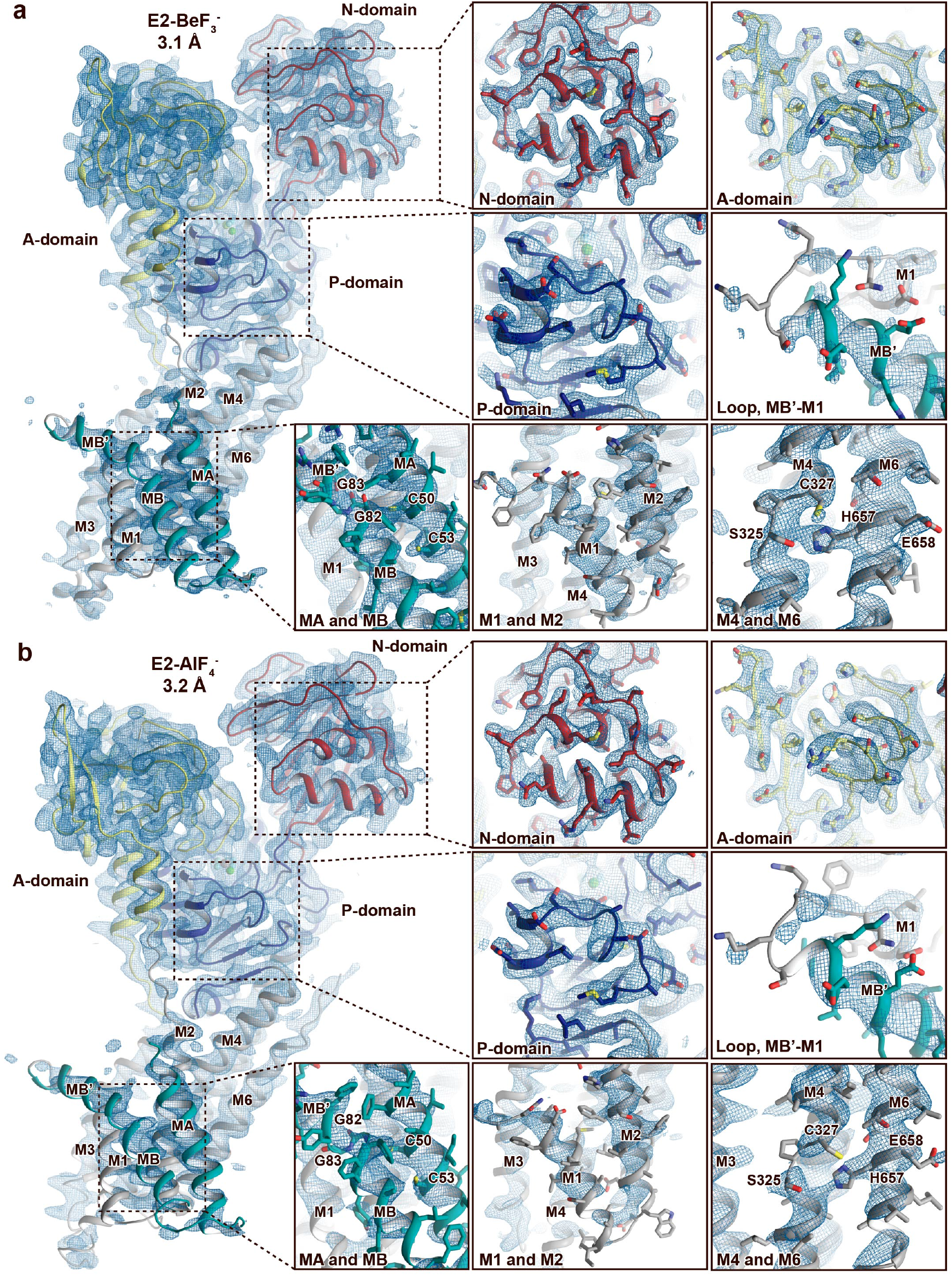
Electron density quality. Final, sharpened, 2Fo-Fc electron density (σ=1.0, blue mesh). The overall resolution is indicated and the structures are colored as in Figure 1. **a**, E2-BeF_3-_ and **b**, E2-AlF_4-_. The quality of the maps differs between structures and domains. The A-, P- and N-domains are well-resolved in both structures. The M-domain is in general less well-resolved than the soluble domains, and the domain is somewhat more well-resolved in the E2-BeF_3-_structure than in E2-AlF_4-_ structure. Nevertheless, it is still clear that MA and MB is present and that C50 and C53 in the N-terminus is membrane embedded and not part of a heavy metal binding domain (HMBD). This is relevant, as CXXC is otherwise a pattern typically linked to a solvent-exposed metal binding site in HMBDs of other P_IB_-groups.

**Supplementary Figure 5.**
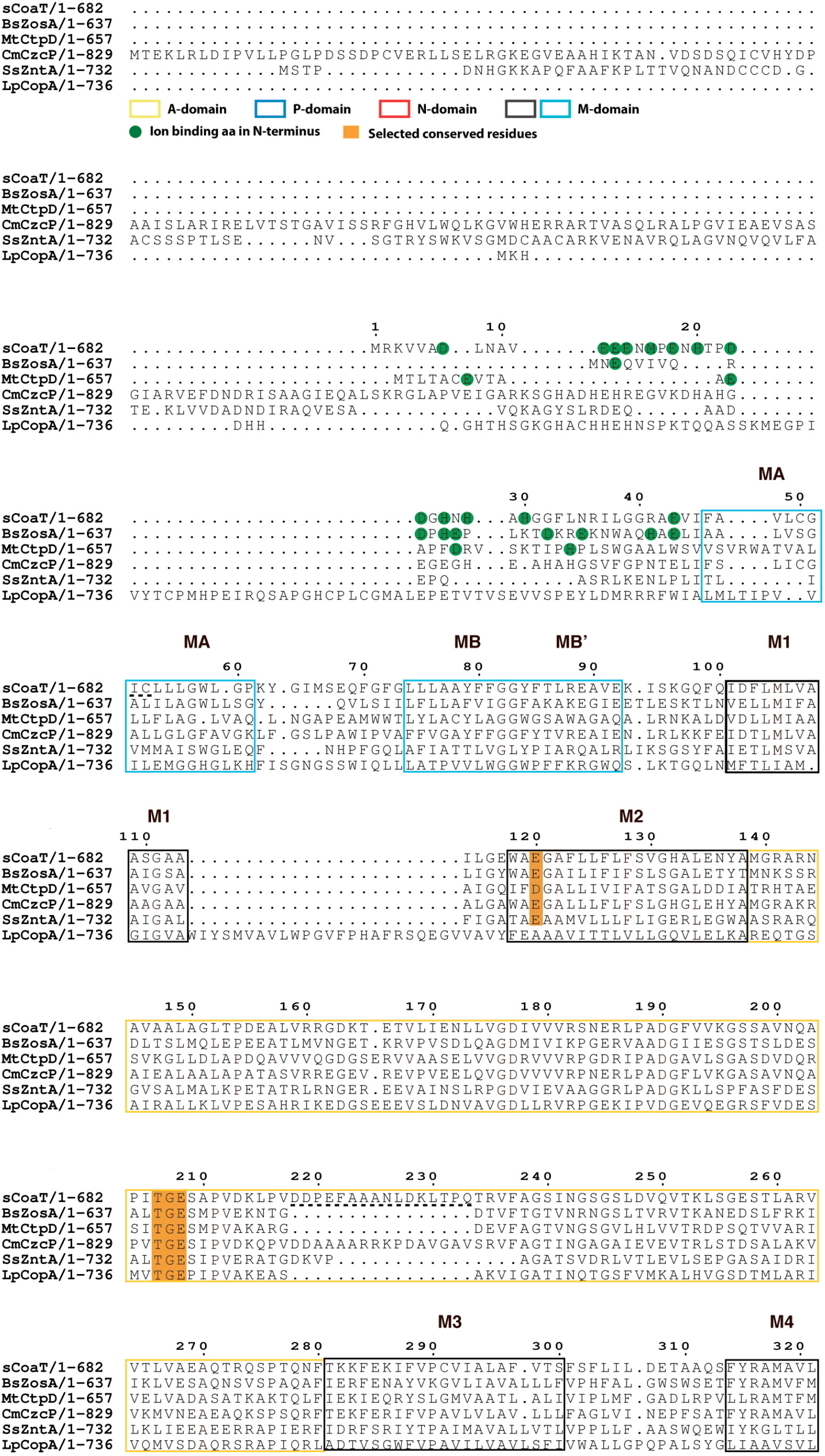
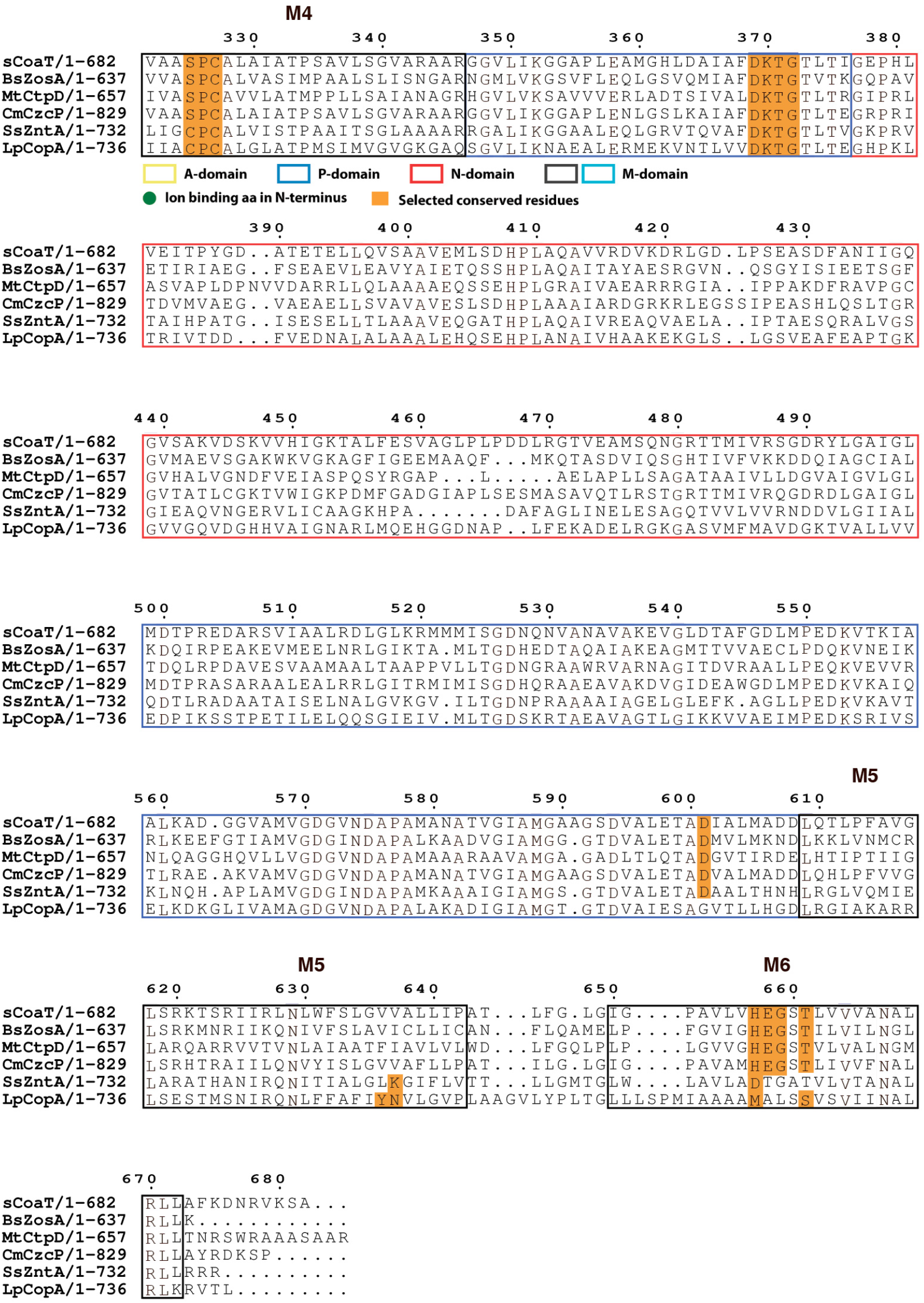
Sequence alignment of selected P_IB_-ATPases. Sequence alignment of four P_IB-4_-ATPases, sCoaT from *Sulfitobacter* sp. NAS-14.1 CmCzcP from *Cupriavidus metallidurans*, BsZoa from *Bacillus subtilis* and MtCtpD from *Mycobacterium tuberculosis*. The P_IB-1_-ATPase LpCopA from *Legionella pneumophila* and the P_IB-2_-ATPase SsZntA from *Shigella sonnei* are also included for comparison.

**Supplementary Figure 6.**
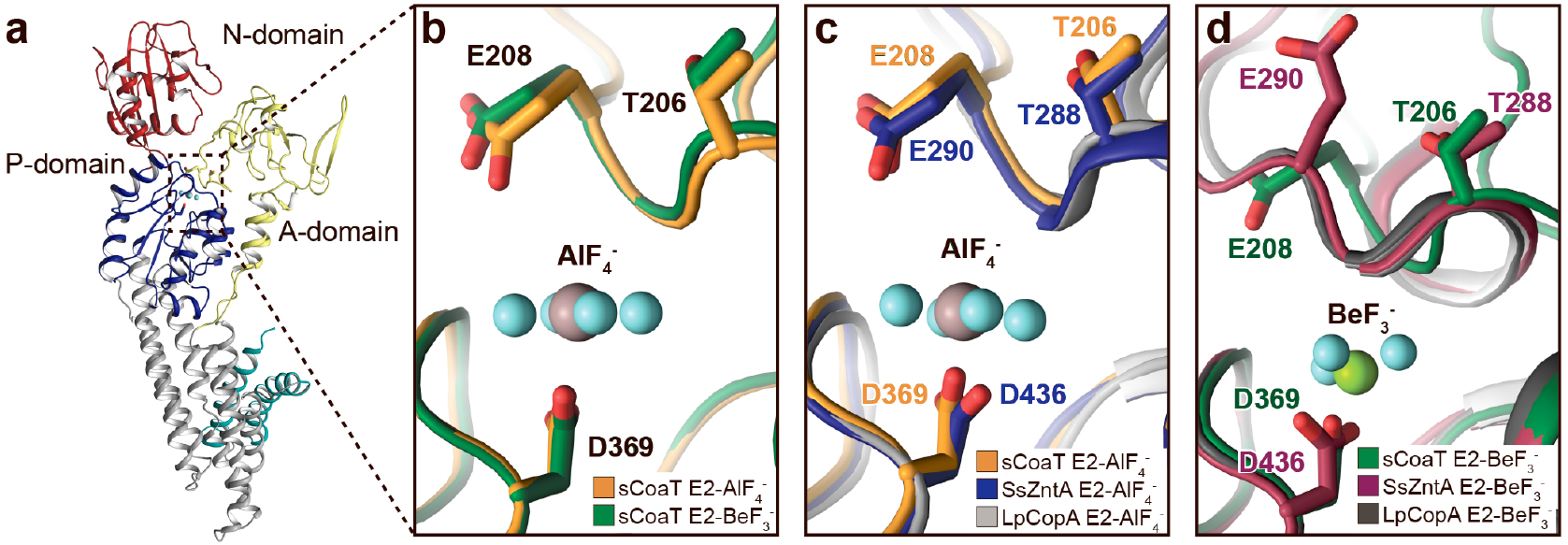
Close view of the phosphorylation site. The TGE loop in the E2-BeF_3-_ stabilized sCoaT (E2P*) is pre-organised for dephosphorylation, which is not the case for SsZntA and LpCopA. **a**, The overall E2P* structure of sCoaT showing the region of focus in panels **b-d. b**, Comparison of the TGE loop in the two sCoaT structures, with only minor differences. **c**, Comparison of sCoaT E2.P_i_ with the equivalent structures of SsZntA and LpCopA (PDB-ID: 4UMW and 4BYG). **d**, Comparison of sCoaT E2P* with the equivalent structures of SsZntA and LpCopA (PDB-ID: 4UMV and 4BBJ).

**Supplementary Figure 7.**
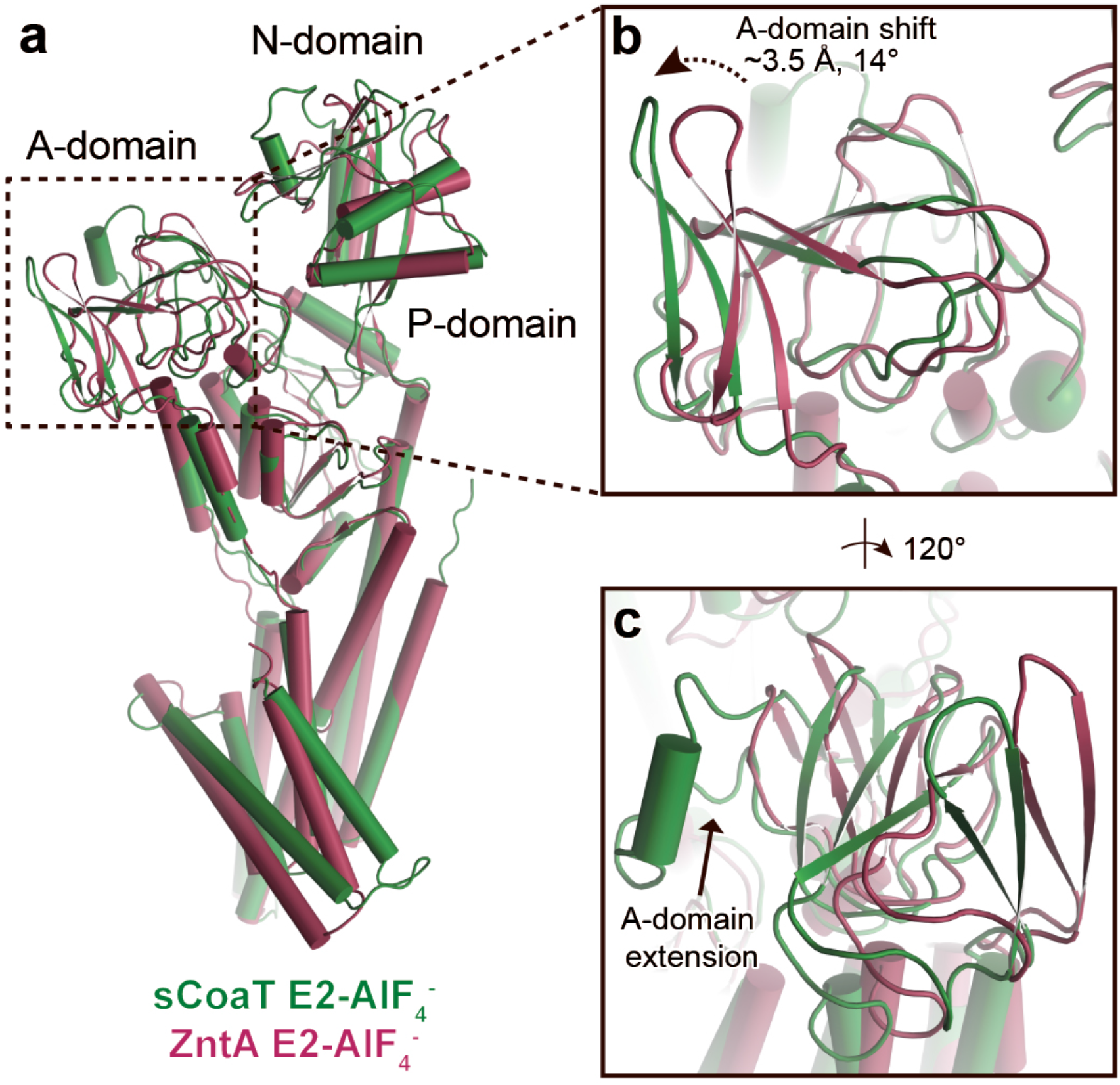
A-domain differences. Superimposition of the E2-AlF_4_ structures of sCoaT (determined here) and SsZntA (PDB ID 4UMW). **a**, The overall structures showing the region of focus in panels **b** and **c. b**, Comparison of the A-domain of sCoaT and SsZntA shows that the peripheral part of the A-domain in sCoaT is shifted closer to the P-domain, whereas the area around the conserved TGE motif (the Glu of the TGE motif is visualized as a sphere) superposes well with SsZntA. **c**, Similar to SERCA, the A-domain of sCoaT possesses a surface-exposed extension which is however not present in P_IB-1_- and P_IB-2_-ATPases.

**Supplementary Figure 8.**
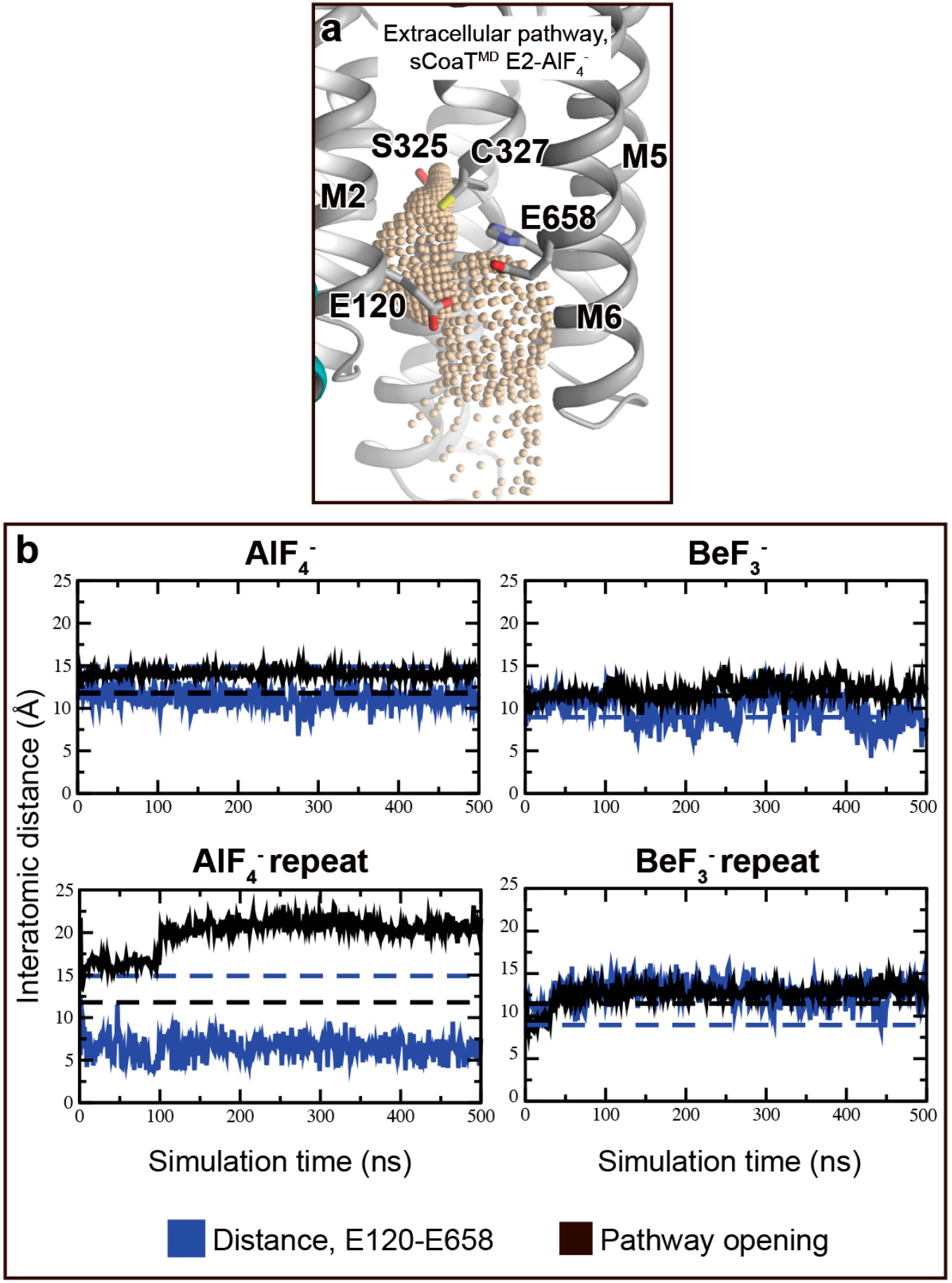
Conformational dynamics during the MD simulations. **a,** The release pathway (wheat) detected in MD simulations of the E2-AlF_4-_ structure. Calculated using the software HOLE. **b**, The E120(oxygen)-E658(oxygen) interatomic distance and the exit pathway opening (W118(Cα)-P652(Cα)) during MD simulations are displayed in blue and black respectively. The corresponding crystal structure distances are shown as dashed lines. The E120-E658 distance decreases in three simulations, with a direct contact formed in the AlF_4-_-repeat simulation. The W118-P652 distance tracks the M2-M6 opening, which is increased in all simulations compared to the crystal structures. Interestingly, the largest opening is achieved in the AlF_4-_-repeat simulation, which also displays the direct E120-E658 interaction, indicating a putative synchronized ion-release mechanism.

## Notes

### Competing Interest Statement

The authors have declared no competing interest.

